# Synapsin is required for dense core vesicle capture and cAMP-dependent neuropeptide release

**DOI:** 10.1101/838953

**Authors:** Szi-chieh Yu, Wagner Steuer Costa, Jana F. Liewald, Jiajie Shao, Alexander Gottschalk

**Affiliations:** Buchmann Institute for Molecular Life Sciences, Goethe-University, Max-von-Laue-Strasse 15, D-60438 Frankfurt, Germany; Department of Biochemistry, Chemistry and Pharmacy, Institute for Biophysical Chemistry, Goethe-University, Max-von-Laue-Strasse 9, D-60438 Frankfurt, Germany

## Abstract

Release of neuropeptides from dense core vesicles (DCVs) is important for neuromodulation. By optogenetics, behavioral analysis, electrophysiology, and electron microscopy, we show that synapsin SNN-1 is required for cAMP-dependent neuropeptide release in *Caenorhabditis elegans* cholinergic motor neurons. In synapsin mutants, behaviors induced by the photoactivated adenylyl cyclase bPAC, which we previously showed to depend on acetylcholine and neuropeptides, are altered like in animals with reduced cAMP. While synapsin mutants have slight alterations in synaptic vesicle distribution, DCVs were affected much more: DCVs were ~30% reduced in synaptic terminals, and not released following bPAC stimulation. Imaging axonal DCV trafficking, also in genome-engineered mutants in the serine-9 protein kinase A phosphorylation site, showed that synapsin captures DCVs at synapses, making them available for release. In non-phosphorylatable SNN-1B(S9A) mutants, DCVs traffic less and accumulate, likely by enhanced tethering to the actin cytoskeleton. Our work establishes synapsin as a key mediator of neuropeptide release.

## INTRODUCTION

Neurotransmitters are released from synaptic vesicles (SVs) by regulated fusion with the plasma membrane (Burkhardt, 2015; Jahn and Fasshauer, 2012; Sudhof, 2012, 2013). Following biogenesis, SVs are filled with neurotransmitter and stored in the reserve pool (RP), by tethering them to the cytoskeleton and to each other, mediated by the phosphoprotein synapsin (Benfenati et al., 1989a; Benfenati et al., 1989b; Cesca et al., 2010; Milovanovic et al., 2018; Rizzoli, 2014; Rizzoli and Betz, 2005). Prior to fusion, SVs are mobilized from the RP by phosphorylation of synapsin by protein kinase A (PKA) and Ca^2+^-calmodulin dependent kinase II (CaMKII) (Kuromi and Kidokoro, 2000; Menegon et al., 2006; Milovanovic et al., 2018), translocated to the plasma membrane (PM), and then become docked and primed by interactions with the proteinaceous scaffold of the active zone, the dense projection (DP) (Sigrist and Schmitz, 2011; Siksou et al., 2007; Stigloher et al., 2011). Primed SVs (forming the readily releasable pool - RRP) fuse with the PM following Ca^2+^ influx into the synaptic terminal. SVs are recycled after endocytosis from endosomal compartments (Haucke et al., 2011; Hua et al., 2013; Jahne et al., 2015; Kittelmann et al., 2013; Kononenko and Haucke, 2015; Rizzoli, 2014; Steuer Costa et al., 2017; Watanabe et al., 2013a; Watanabe et al., 2013b; Watanabe et al., 2014; Yu et al., 2018).

The rates of SV mobilization and priming need to adapt to the neurons’ activity, i.e. the frequency of SV fusion events. Thus, mechanisms ruling the rates of recycling, mobilization and priming of SVs need to be in place, but also quantal size may be modulated (Liu et al., 2005; Steuer Costa et al., 2017; Wierda et al., 2007; Xue and Wu, 2010). Such mechanisms depend on intra- and extracellular Ca^2+^ concentrations (Liu et al., 2005; Schneggenburger and Rosenmund, 2015). During sustained neuronal activity, elevated cytosolic Ca^2+^ may further induce secretion of DCVs via protein kinase C (PKC) signaling (Bark et al., 2012; Park et al., 2006; Sieburth et al., 2007; Xue and Wu, 2010). An additional pathway of presynaptic modulation is regulated by cAMP and PKA (Baba et al., 2005; Gekel and Neher, 2008; Wang and Sieburth, 2013). This affects SV mobilization and thus their release, however, an unknown PKA target was found to also affect DCV fusion (Zhou et al., 2007). PKA modulates several synaptic proteins by phosphorylation, e.g. synapsin (Hosaka et al., 1999), tomosyn (Baba et al., 2005; Gracheva et al., 2006; McEwen et al., 2006), Rim1 (Lonart et al., 2003), the ryanodine receptor (RyR; (Rodriguez et al., 2003), cysteine string protein (Evans et al., 2001), Snapin (Thakur et al., 2004), complexin (Cho et al., 2015), and SNAP-25 (Nagy et al., 2004). Likewise, Exchange Protein Activated by Cyclic AMP (EPAC) regulates SV release in *Drosophila*, crayfish and rats (Cheung et al., 2006; Gekel and Neher, 2008; Kaneko and Takahashi, 2004; Zhong and Zucker, 2005).

Neuropeptides regulate a plethora of processes in the nervous system and control alteration of brain states (Graebner et al., 2015; Oranth et al., 2018; Steuer Costa et al., 2019; Taghert and Nitabach, 2012). They are synthesized as precursor proteins, and packaged and processed in DCVs while being transported to neuronal terminals (Hoover et al., 2014). In *Drosophila*, constant antero- and retrograde DCV trafficking occurs between synapses, and capture events at terminals make DCVs available for release (Wong et al., 2012). Little is known about proteins mediating DCV capture, however, the process appears to be regulated in an activity-dependent fashion, and was shown to involve fragile-X mental retardation-like protein (FMRP) in flies (Cavolo et al., 2016), as well as synaptotagmin 4. The latter protein was more involved in regulating the interaction of DCV cargo with the microtubule-linked motor proteins kinesin and dynein, thus capture may depend on a tug-of-war between these antero- and retrograde motors (Bharat et al., 2017). However, also actin was found to play a role, such that localization of DCVs at neuronal terminals may involve a handover from microtubules to the actin cytoskeleton. In line with this, a requirement for the scaffold proteins α-liprin and TANC2 was shown in DCV capture at postsynaptic spines, and these proteins interact with actin as well to modulate function of kinesin motors, which also involves calmodulin (Stucchi et al., 2018). Last, in *C. elegans*, a complex of proteins involving sentryn, α-liprin and the SAD kinase were shown to regulate antero- vs. retrograde traffic of DCVs, thus ‘capture’ could be determined by a balance of the two processes (Morrison et al., 2018). Whether there are proteins that physically attach DCVs to the cytoskeleton near release sites is not understood.

Release of neuropeptides from DCVs is modulated by cAMP/PKA as well as PKC signaling (Sieburth et al., 2007; Steuer Costa et al., 2017), and requires the Ca^2+^-dependent activator protein for secretion (CAPS/UNC-31) (Charlie et al., 2006a; Hammarlund et al., 2008; Rupnik et al., 2000; Speese et al., 2007; Steuer Costa et al., 2019). We have shown that optogenetic cAMP increase, induced by *Beggiatoa sp.* photoactivated adenylyl cyclase (bPAC), causes fusion of DCVs in *C. elegans* cholinergic motor neurons (Steuer Costa et al., 2017). This resulted in exaggerated yet coordinated movement, indicating that ACh output was enhanced, but that the stimulation did not override intrinsic activity patterns. bPAC stimulation caused a higher frequency of miniature postsynaptic current (mPSC) events, which represent fusion of single or few SVs. Interestingly, bPAC, but not ChR2, stimulation led to an increase in mPSC amplitudes, which required UNC-31/CAPS, thus neuropeptide signaling was involved. These neuropeptides were released from cholinergic neurons, acting in an autocrine fashion. We could show that the neuropeptides induced additional filling of SVs with ACh via the vAChT (UNC-17 in *C. elegans*), evident also at the ultrastructural level, as an increase in SV diameter. cAMP affected behavioral changes not only by increased SV loading, but also by causing a higher SV release rate, as well as by increased SV mobilization from the reserve pool.

Here, we identify synapsin as a main mediator of cAMP/PKA effects on synaptic transmission, DCV trafficking and fusion. Synapsin *snn-1* mutants did not release any neuropeptides and accumulated them in neuronal cell bodies and processes. *snn-1* mutants also exhibited reduced release of SVs, which distributed abnormally, in line with SNN-1 organizing the RRP, and SVs did not show the bPAC evoked, neuropeptide dependent increase in diameter and ACh content. Consistent with a defect in DCV release, DCV numbers were reduced in *snn-1* mutants and their distribution in synaptic terminals differed from wild type, indicating a defect of *snn-1* synapses to capture DCVs and to retain them for release. Trafficking of DCVs was aberrant in *snn-1* deletion mutants and particularly affected in SNN-1(S9A) knock-in animals, with non-phosphorylatable SNN-1B, where DCVs showed strongly reduced motility, likely by enhanced tethering of DCVs to the actin cytoskeleton via SNN-1.

## RESULTS

The augmenting effects of cAMP on synaptic transmission involve PKA signaling. To further clarify this pathway, we wanted to identify the target(s) of cAMP/PKA that mediate the effects on SV mobilization and release, and also on the fusion of neuropeptide containing DCVs. As we found previously, bPAC stimulation in cholinergic neurons increased locomotion speed and body bending angles (Steuer Costa et al., 2017), thus we quantified these behavioral parameters to analyze various mutants of candidate synaptic cAMP and PKA targets for potential alteration in these bPAC-evoked phenotypes. A directly cAMP-regulated modulator of synaptic strength is EPAC (encoded by *epac-1*; Kaneko and Takahashi, 2004). We focused on the following further proteins that are PKA targets: The UNC-68 ryanodine receptor (RyR), a major mediator of intracellular Ca^2+^ signals, absence of which affects synaptic signaling (decreased mPSCs frequency, lack of high amplitude mPSCs; Liu et al., 2005). The single *C. elegans* synapsin (*snn-1)* resembles vertebrate synapsin II, which maintains the SV reserve pool by tethering SVs to the cytoskeleton (Benfenati et al., 1989a; Gitler et al., 2008). PKA phosphorylation of synapsin mobilizes SVs from the reserve pool (Menegon et al., 2006) by releasing their mutual and cytoskeletal anchoring, thus promoting their mobilization to the RRP (Hosaka et al., 1999; Johnson et al., 1972). More recent work showed that the C-terminus of synapsin contains an intrinsic disordered region (IDR) that can form non-membranous organelles by liquid-liquid phase separation in a phosphorylation state dependent manner, regulated by activity and CaMKII (Milovanovic et al., 2018). As positive controls, we also included *unc-31* (CAPS) mutants (lacking neuropeptide release) and PDE(gf) animals expressing a constitutively active phosphodiesterase (Charlie et al., 2006b), with reduced cAMP levels. We plotted the change of locomotion bending angles or speed, relative to before illumination (**Fig. 1a-d**). To compare the different genotypes and conditions for similar phenotypes, which may point to similar mechanisms underlying the effects on the induced behaviors, we used dynamic cluster analysis.

**Fig. 1:**
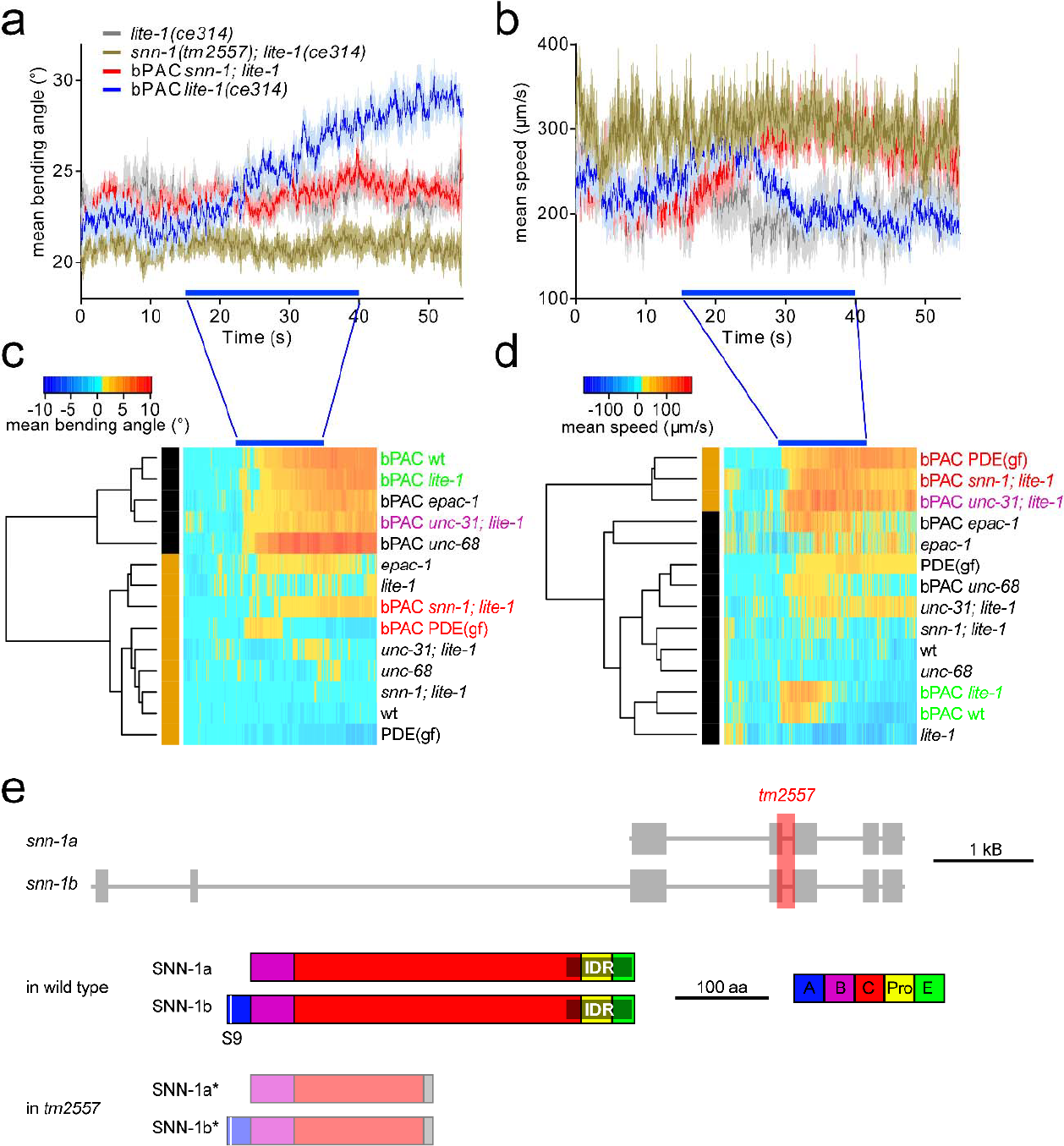
Behavioral phenotypes induced by bPAC photostimulation in cholinergic neurons uncover synapsin as a main target of cAMP increase. bPAC was expressed in cholinergic neurons and light-effects on locomotion behavior analyzed. **a)** Mean (± SEM) bending angles (n ≥ 29), measured for 11 angles defined by 13 points along the body or **b)** velocity of animals before, during and after blue light stimulation (blue bar). Animals of the indicated genotypes were assessed. **c)** Bending angles and **d)** velocities for n ≥ 29 animals of the indicated genotypes and transgenes were compared by dynamic cluster analysis, following normalization to the mean before light stimulation. Two clusters are observed for both behaviors, one with wild type animals (green), and another with a gain-of-function phosphodiesterase (PDE(gf)) and *snn-1* mutants (red). **e)** Gene locus of *snn-1* in *C. elegans*, on chromosome IV (boxes, exons, lines, introns), as well as protein structure of the a and b splice variants of SNN-1. Domain nomenclature A, B, C, Pro (proline-rich), E, according to homology to the mammalian isoforms (Sudhof et al., 1989). Domains A and E are required for SV and synapsin oligomerization, while domain C is required for interaction of synapsin with the SV membrane. The intrinsic disordered region (IDR, grey shade) in the C-terminal half, recently shown to mediate formation of phase separated condensates of synapsin and synaptic vesicles (Milovanovic et al., 2018), was annotated based on a prediction using SNN-1B, and the ‘Predictor of Natural Disordered Regions’ web interface (www.pondr.com). S9 denotes the main phosphorylation site for PKA, however, numerous phosphorylation sites for different kinases exist all over the protein. For review, see (Cesca et al., 2010).

bPAC-expressing animals were assessed in wild type (wt) or in *lite-1* background, to avoid non-specific behavioral responses to blue light (LITE-1 is a photosensor that mediates intrinsic UV-/blue-light avoidance of *C. elegans*; Edwards et al., 2008). Upon photostimulation, bPAC expressing animals showed a progressive increase of bending angles (**Fig. 1a, c**). This was also observed in *epac-1*, *unc-68* and *unc-31* mutants, but not in *snn-1* and PDE(gf) animals. bPAC stimulated animals also showed a moderate speed increase, which was similar in *unc-68(r1162)* and *epac-1(ok655)* mutants expressing bPAC. Yet, a much larger speed increase was observed for *unc-31(n1304)*, PDE(gf) and *snn-1(tm2557)* mutants (**Fig. 1b, d**). The *snn-1(tm2557)* allele truncates the 3’-half of the gene, likely eliminating its expression (**Fig. 1e**). Even if some expression remains, only the N-terminal half, including the PKA phosphorylation site I (serine 9), would be produced, while the truncation of the central C domain, interacting with SVs and actin, would abolish synapsin function. *snn-1(tm2557)* mutants were the only animals that mimicked PDE(gf) mutants in both behaviors, differing from wt, and they phenocopied *unc-31* mutants in speed increase. We therefore analyzed *snn-1* mutants in more detail.

### Synapsin is required for neuropeptide release

First, we assessed bPAC-induced neuropeptide release. When neuropeptide precursors are fused to a fluorescent protein, their exocytosis can be detected in so-called scavenger cells (coelomocytes). These have kidney-like function and ‘clean up’ the body fluid by endocytosis. Coelomocytes become fluorescent, when Venus-tagged NLP-21 neuropeptides are expressed in motor neurons (NLP-21 is expressed intrinsically in these cells; Sieburth et al., 2007; **Fig. 2a**). bPAC stimulation induced NLP-21∷Venus release, as we showed previously (Steuer Costa et al., 2017). However, this was abolished in *snn-1(tm2557)* mutants (**Fig. 2b, c**), despite a significant enrichment of NLP-21∷Venus in neuronal cell bodies and processes, compared to wt (**Fig. 2d, e**). The latter may result from an inability of *snn-1* mutants to release neuropeptides.

**Fig. 2:**
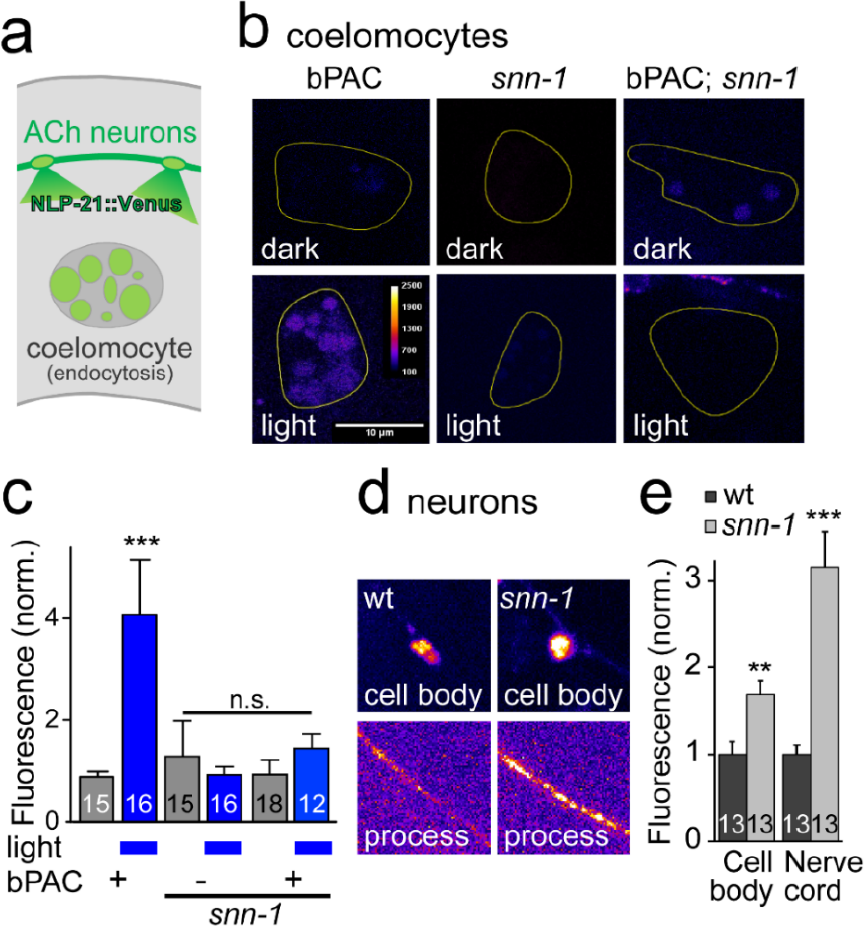
Synapsin is required for cAMP-induced neuropeptide release. **a)** NLP-21∷Venus neuropeptides are released from cholinergic neurons and endocytosed by coelomocytes. **b)** Representative images of coelomocytes in bPAC, *snn-1* and *snn-1*; bPAC animals, without and with photostimulation. False color representation, scale bar 10 μm. **c)** Fluorescence quantification in coelomocytes, normalized to fluorescence level in wt, non-stimulated. **d)** Fluorescence and **e)** quantification of NLP-21∷Venus in cell bodies and processes of cholinergic neurons, in wt and *snn-1(tm2557)* mutants. Number of animals indicated. Mean ± SEM, in c) and e), statistically significant differences (**p<0.01; ***p<0.001) were determined by one-way ANOVA with Bonferroni correction.

Previously, we demonstrated that bPAC-stimulation increases the SV release rate, and the amplitude of mPSCs in patch-clamp recordings (Steuer Costa et al., 2017). The amplitude increase resulted from neuropeptide release, impacting on filling of SVs with ACh, and mutation of *unc-31* abolished the mPSC amplitude increase. In *snn-1(tm2557)* mutants, the basal mPSC rate, on average, was slightly reduced (**Fig. 3a-b**), though this was not significantly different from wt (**Fig. 3d**). bPAC stimulation increased the mPSC rate in wt, and likewise in *snn-1(tm2557)* animals (**Fig. 3a, b, d**). Yet, like *unc-31* mutants, *snn-1* animals showed no increased mPSC amplitudes (**Fig. 3c, e**). This suggests that synapsin may be required for neuropeptide release.

**Fig. 3:**
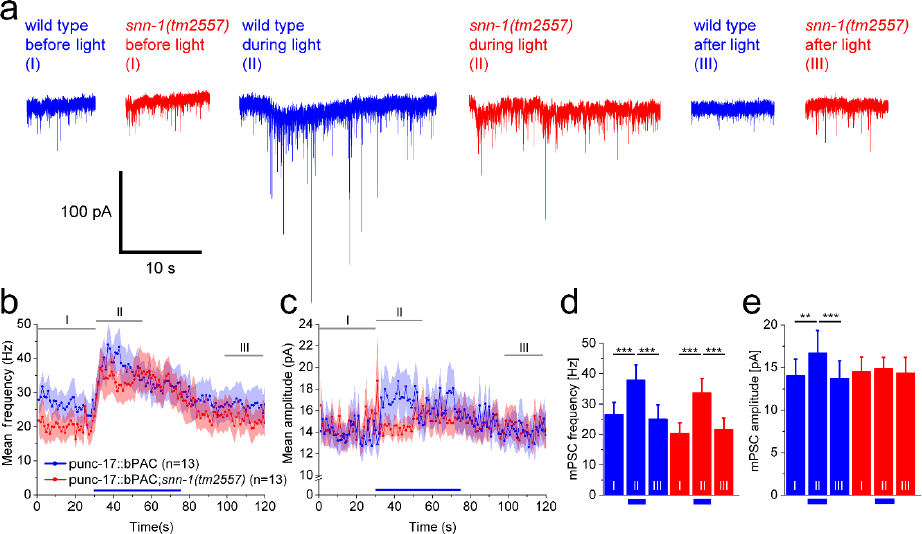
Postsynaptic currents at the neuromuscular junction (NMJ) after presynaptic bPAC stimulation uncover SNN-1 function in cAMP-dependent amplitude increase. **a)** Postsynaptic currents (voltage clamp of body wall muscle cells) before (I), during (II) and after (III) presynaptic bPAC stimulation in cholinergic neurons, in wt (blue) or *snn-1(tm2557)* background (red). **b)** Mean (± SEM) mPSC events per second and **c)** mPSC amplitudes, in 1 s bins, before, during and after photoactivation (blue bar), in animals expressing bPAC in wt and *snn-1* mutants. **d, e)** group data from b, c), averaged in the indicated time windows I, II, III. Statistically significant differences (**p<0.01; ***p<0.001) were determined by paired students t-test.

### Synapsin mutants have normal SV numbers, but reduced DCV numbers at synapses

To assess the role of synapsin in synaptic transmission, and specifically in neuropeptide release, we turned to ultrastructural analysis. High-pressure-freeze electron microscopy (HPF-EM; Kittelmann et al., 2013; Yu et al., 2018) can be used to analyze the content and distribution of synaptic organelles in the terminals of cholinergic motor neurons, including SVs, docked SVs, DCVs and large vesicles (LVs), which are endosomes formed after activity-induced SV release (**Fig. 4a, b**).

**Fig. 4:**
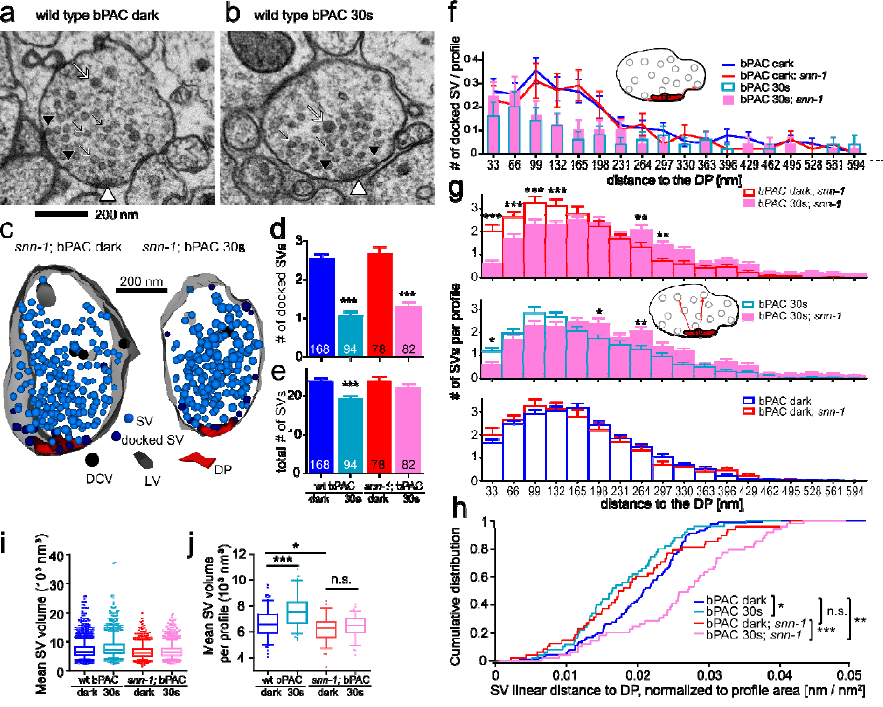
Electron microscopy analysis of the NMJ reveals reduced SV release, altered SV localization and smaller SV diameter in *snn-1(tm2557)* mutants. **a)** Thin section of a wt cholinergic neuron synaptic terminal. White arrowhead points to dense projection (DP). Black arrows: synaptic vesicles (SVs); black arrowheads: docked synaptic vesicles; white arrow: dense core vesicle (DCVs). **b)** As in a), but after 30 s blue light stimulation. **c)** 3D-reconstruction of *snn-*1; bPAC synapses, dark control (left) and 30 s photostimulated (right). Synaptic structures, as in a, b) are indicated, including large vesicles (LVs) that result from endocytosis following SV fusion. **d)** Number of docked SVs and **e)** total SVs per profile, normalized to perimeter or area, respectively. **f)** Binned distribution of docked SVs per profile, at indicated distances (along the plasma membrane) to DP, in wt or *snn-1(tm2557)* before and after 30 s stimulation. **g)** Binned distribution of SVs in the reserve pool in *snn-1* versus wt, before and after 30 s stimulation, at indicated distances to the DP. **h)** Empirical cumulative distribution plot of summed distances of reserve pool SVs to DP per profile. **i)** Volume of SVs (n=4,067; 2,079; 1,400 and 1,888), for the indicated genotypes and conditions of bPAC stimulated synapses. SV inner diameter was measured and used to calculate the volume. Also shown are means, medians, interquartile range, whiskers: 5-95 percentile). All statistical comparisons (ANOVA with Bonferroni correction) were highly significant, due to the high n numbers. **j)** SV volumes, as in i), were averaged per profile, for profiles with ≥10 SVs (n=168, 94, 78, and 82). Shown are means±SEM, *p≤0.05, **p≤0.01, ***≤0.001, one-way ANOVA with Tukey’s multiple comparison test in d, e, g, j and Bonferroni correction in i; Kolmogorov-Smirnov test in h.

First, we analyzed the distribution of SVs. We had previously shown that bPAC stimulation strongly reduced the number of docked SVs, as well as overall SV numbers throughout the terminal (i.e., the sum of RRP and reserve pool) in wild type animals (Steuer Costa et al., 2017). We now compared this in *snn-1(tm2557)* mutants (**Fig. 4c-e**). Like in wt, bPAC stimulation caused significantly reduced docked SVs in *snn-1* mutants. However, unlike in wt, no significant reduction of the reserve pool was observed in *snn-1* animals (**Fig. 4d, e**). Distribution of docked SVs, relative to the dense projection (DP; central to the active zone - AZ) was similarly altered by bPAC stimulation in *snn-1* and wt (**Fig. 4f**). Synapsin therefore does not directly affect SV priming or exocytosis. However, as the mPSC rate was somewhat reduced in *snn-1* animals, it could facilitate these processes by promoting SV transfer from the RP to the RRP.

Analysis of the RP distribution showed that SVs which locate near the DP during bPAC photostimulation in wt were depleted from the DP vicinity in *snn-1* (**Fig. 4g, h**). If SVs approach the AZ by tethering to the DP, and then are docked and distributed laterally, depletion of SVs near the DP may reflect a deficit in refilling of the RRP of docked SVs. Thus, our finding supports a role of synapsin in SV tethering and mobilization from the RP. Compared to wt synapses, *snn-1* synapses were smaller (**Fig. S1a, b**) and did not accumulate endocytic large vesicles upon bPAC stimulation (**Fig. S1e**), probably as reduced SV release necessitated less SV recycling. Nonetheless, *snn-1* synapses increased their size in response to 30 s bPAC stimulation, as did wt. Finally, we also analyzed the SV diameter in *snn-1* mutants, since we previously found that increased SV loading due to cAMP and neuropeptide signaling led to larger SVs (Steuer Costa et al., 2017). In comparison to wt, the *snn-1* mutants had significantly smaller SVs, which did not increase their size upon bPAC stimulation (**Fig. 4i, j**), as we previously found for *unc-31* mutants (Steuer Costa et al., 2017). This indicates that, in line with our findings of abolished neuropeptide release, *snn-1* mutants also do not upregulate SV filling. It further indicates that SV size is actively regulated by signaling via as yet unknown neuropeptides.

Second, we analyzed the distribution and fate of DCVs in *snn-1(tm2557)* synapses before and after bPAC-stimulation. DCVs were significantly reduced in *snn-1* mutants, compared to wt, under both conditions, in sections containing the DP (**Fig. 5a, b**), and in sections neighboring the DP (to ~240 nm axial DP distance; **Fig. 5c, d, f**). We also assessed the radial distribution of DCVs relative to the DP. DCVs were most abundant in 150-250 nm distance to the DP, peaking at about 233 nm (**Fig. 5d, e**). Although bPAC stimulation caused DCV depletion in the 150 nm bin (**Fig. 5e**), overall distribution and abundance were similar to without stimulation, peaking at 200-230 nm radial DP distance in wt. However, in *snn-1(tm2557)* mutants, DCVs peaked at 250-350 nm radial distance, with no marked further depletion upon bPAC stimulation. The reduction of DCVs in the *snn-1* mutant indicates that DCVs are delivered to *snn-1* terminals, but may not be efficiently tethered there to be eventually released. Thus, they may be ‘lost’ to the neuronal process and cell body (**Fig. 2d**). As reported previously (Hammarlund et al., 2008), DCVs did not cluster at AZs, but evenly distributed along the axon. DCVs in sections flanking the DP out to 240 nm (**Fig. 5f**) did not exhibit any obvious peak, yet they were depleted in *snn-1* synapses. In sum, *snn-1* mutants have reduced DCV numbers in synapses, which are not released in response to bPAC stimulation.

**Fig. 5:**
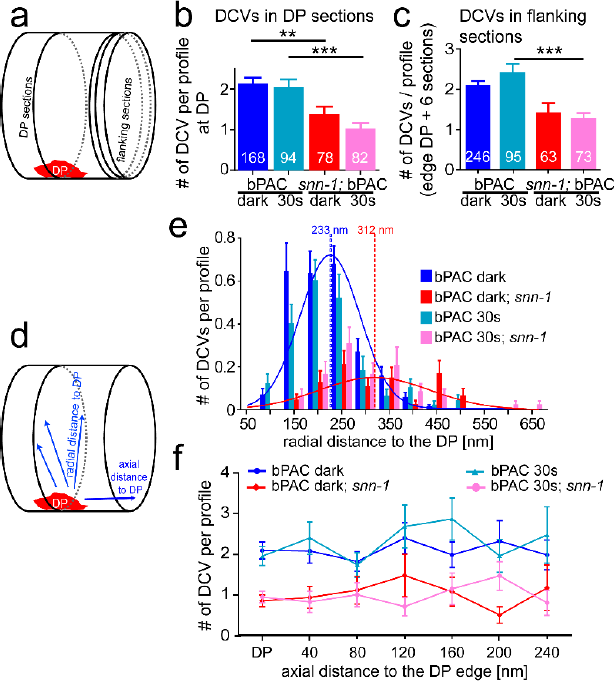
DCVs are largely depleted in *snn-1(tm2557)* synapses and distribute differently than in wt. **a)** Sections analyzed by HPF-EM either contain the DP or are flanking the region containing the DP. **b)** DCVs per profile containing the DP in *snn-1(tm2557)* compared to wt, without and with 30 s photo-stimulation. **c)** Same as in b), but in DP-flanking sections. **d)** Abundance of DCVs was analyzed either radially within a section, in distinct distances to the DP, or along the axon. **e)** Abundance of DCVs in distinct radial distances of DCVs relative to DP, quantified in 50 nm bins in untreated and 30 s stimulated wt and *snn-1(tm2557)* synapses. **f)** DCV abundance in sections containing the DP, or in sections of the indicated axial distance to the DP. b, c, e, f) Mean±SEM. *p≤0.05, **p≤0.01, ***≤0.001, one-way ANOVA with Tukey correction.

### Synapsin S9A and S9E phosphorylation site mutants show behavioral and electrophysiological defects

As we showed previously (Steuer Costa et al., 2017), cAMP augments SV release and induces neuropeptide release, and here we demonstrated that this is facilitated by synapsin. Despite effects on distribution and mobilization, the bPAC-induced increase of SV release was not abolished in *snn-1(tm2557)* mutants, which may retain the N-terminal half of the protein (**Fig. 1e**). PKA-mediated phosphorylation of serine 9 reduces the affinity for actin, and was suggested to regulate SV mobilization (Cesca et al., 2010; Hosaka et al., 1999). To explore this in *C. elegans* and to assess the potential function of the S9 residue in synapsin SNN-1B function, we obtained point mutations by CRISPR/Cas9 mediated genome editing: The mutation S9A abolishes phosphorylation and mimics a constitutively dephosphorylated state, and S9E mimics constitutively phosphorylated synapsin. First, we assessed their behavioral phenotypes (**Fig. 6a-d**). Upon photostimulation, bending angles were increasing in both S9A and S9E mutants, however, at the late phase of the 25 s stimulation period, the two point mutants had significantly higher bending angles than wt, with the most pronounced effect for S9E that showed sustained high bending angles after the stimulus ended. For locomotion speed, the S9A animals showed a higher increase than the wt controls, while S9E animals showed an unaltered speed increase.

**Fig. 6:**
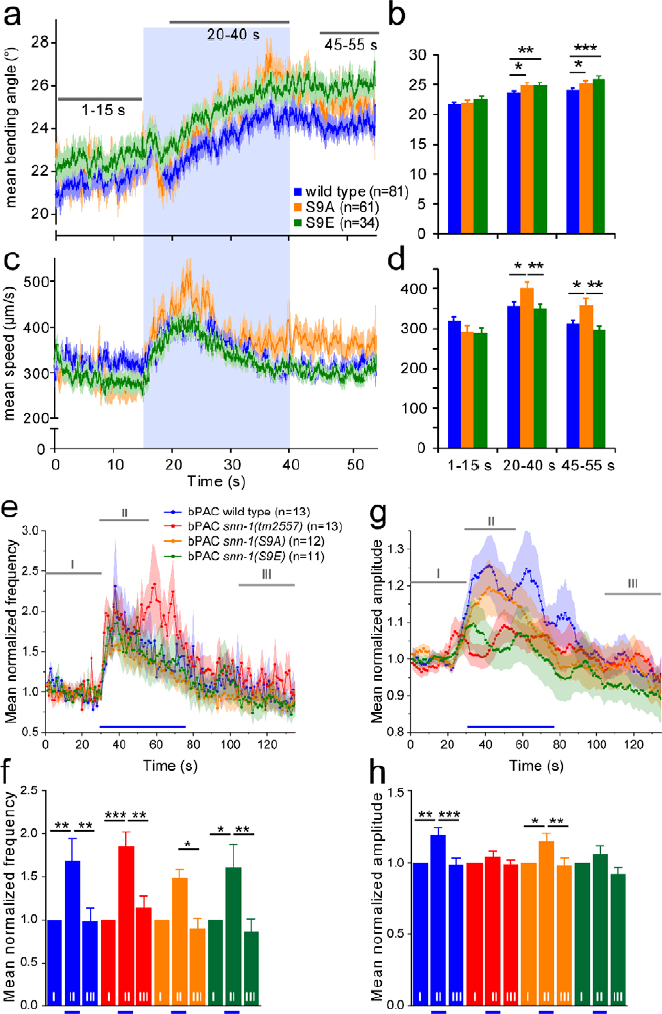
SNN-1B serine 9 mutations affect behaviors and postsynaptic currents induced by bPAC stimulation. SNN-1B S9A and S9E mutations affect bPAC induced changes in crawling bending angles **(a, b)** and velocity **(c, d)** compared to wild type. b, d) Mean (±SEM) bending angles and velocities in the indicated time windows; illumination period indicated by a blue bar in a, c). **e, f)** Mean, normalized mPSC events per second before (I), during (II) and after (III) presynaptic bPAC stimulation in cholinergic neurons, in the indicated genotypes. Blue light stimulation period indicated by blue bar. **g, h)** Mean (± SEM) normalized mPSC amplitudes in 1 s bins, before, during and after photoactivation. Statistically significant differences (*p<0.05; **p<0.01; ***p<0.001) are determined by paired and unpaired t-test (b, d) or two-way ANOVA (f, h) with Bonferroni posttest.

Next, we assessed the synapsin S9 mutants by electrophysiology. Both mutants had essentially the same relative increase in mPSC rate during bPAC stimulation as wt (**Fig. 6e, f**), and as *snn-1(tm2557)* mutants (likely lacking the entire SNN-1B protein; **Fig. 1e**). The latter mutant had a prolonged increase in mPSC rates compared to the other genotypes. When we analyzed the mPSC amplitudes, the S9A mutant showed increased amplitudes, like wt, while the S9E mutants, just as the *snn-1(tm2557)* mutant, showed no amplitude increase during bPAC stimulation (**Fig. 6g, h**). This indicates that SNN-1B(S9E) animals do not release neuropeptides, which could be due to an inability to tether DCVs to the cytoskeleton and to capture them near synaptic release sites.

We assessed this possibility more directly by HPF-EM. Therefore, the axial distribution of DCVs in the S9A and S9E mutants was analyzed to distances ‘out’ to 800 nm from the DP/synapse region (**Fig. 7a-d**). All mutants showed significantly less DCVs than wt, both without and with bPAC stimulation (**Fig. 7e**; **Supplementary Fig. S2**).

**Fig. 7:**
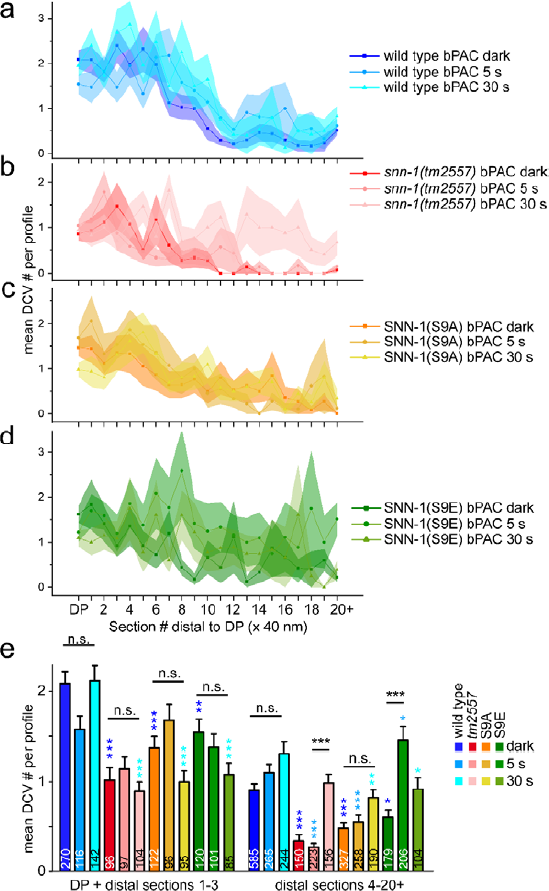
SNN-1B serine 9 mutations affect DCV distribution in sections containing dense projections and flanking sections following bPAC stimulation. DCV distribution and abundance was analyzed in the DP containing and DP flanking sections, out to 800 nm (in 20 consecutive 40 nm sections) in wild type **(a)**, *snn-1(tm2557)* **(b)**, SNN-1B S9A **(c)** and S9E **(d)** knock-in mutants, in unstimulated, as well as 5 s and 30 s bPAC photostimulated animals. Shown are mean±SEM DCV number per section. **e)** Mean (±SEM) DCV numbers were binned in the indicated sections at and near the DP (DP + flanking sections 1-3) or in the distal section (sections 4-20). n=number of sections is indicated in e). Statistical analysis in e) one-way ANOVA with Bonferroni’s multiple comparison test. (*p<0.05, **p<0.01, ***p<0.001). Different blue colored asterisks indicate significance to the respective wild type controls.

In addition, following photostimulation, we found a significant enrichment of DCVs at sites distal to the synaptic terminal / DP region in the *snn-1(tm2557)* and the S9E mutants, while S9A and wt did not increase DCV numbers distal to the DP (**Fig. 7a-e**). This could indicate that the *tm2557* and S9E mutants, either due to the lack of SNN-1, or due to the inability to capture DCVs because of the constitutively ‘phosphorylated’ S9 residue, are unable to accumulate DCVs at synapses. Instead, the few DCVs present there are even further depleted, while DCVs accumulate outside synapses, as was evident also by NLP-21∷Venus imaging (**Fig. 2d**).

### In SNN-1(S9A) mutants, DCVs accumulate and become immobilized in axonal processes

Finally, we analyzed the dynamic properties of DCVs, i.e. their trafficking in the cholinergic motor neurons. We used time lapse imaging of NLP-21∷Venus containing DCVs, expressed in the dorsally innervating A-type (DA) class of motor neurons (**Fig. 8a**; **Supplementary Video S1**). Axonal processes on the dorsal side were imaged as confocal stacks at 4 volumes / s, and maximum projections of the different focal planes ensured that all DCVs were imaged, despite small movement in the focal plane. A line scan through the axon then allowed to generate kymographs, and to analyze antero- as well as retrograde traffic of DCVs, as well as their stationary accumulations (**Fig. 8b**).

**Fig. 8:**
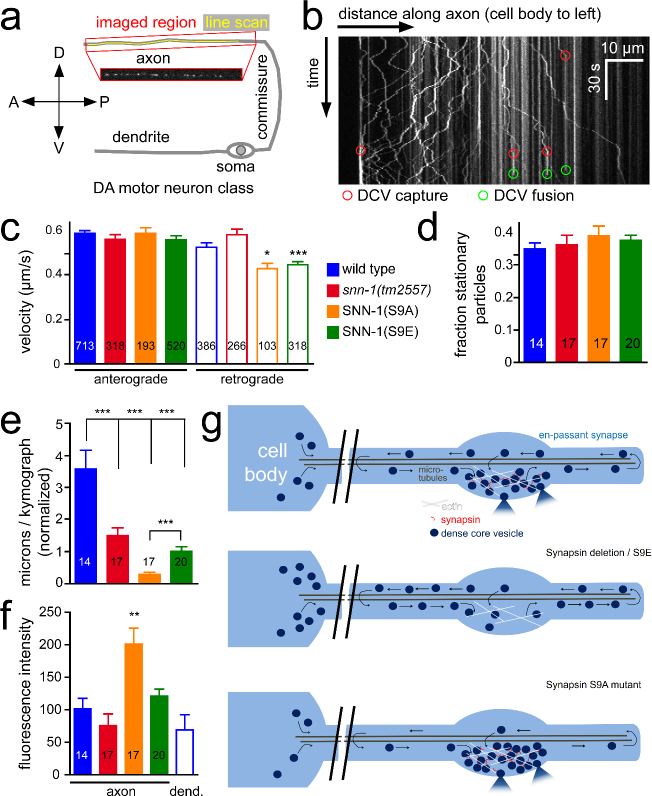
DCV trafficking in axons of DA-type motor neurons is affected in synapsin mutants. DCVs labeled with the NLP-21∷Venus neuropeptide precursor protein were analyzed in axonal processes of DA motor neurons. **a)** Schematic showing the region analyzed and location of the line scan used for kymograph analysis as in b). **b)** Example kymograph, 80 μm along the axon pointing away from the cell body (anterograde traffic is to the right), and covering 105 s in the time domain (down). Trafficking is observable, some DCVs appear to be static. Some putative capture events (red circles) are seen, where a moving particle becomes static. Putative DCV fusion events (disappearance of DCV fluorescence) are evident (green circles). **c)** Velocity (anterograde and retrograde, as indicated) of the DCVs was analyzed in increments of straight movement, between observable changes in velocity, or phases of no movement. Number of analyzed increments is indicated. The analyzed genotypes are indicated. **d)** The fraction of stationary particles was analyzed. **e)** Overall movement was assessed by summing up the individual distances of each moving increment in axial direction. **f)** The density of the fluorescence in the analyzed axonal segments was compared. d-f) Mean±SEM for the indicated number of animals. Statistically significant differences (*p<0.05; **p<0.01; ***p<0.001) are determined by one-way ANOVA with Bonferroni correction. **g)** Model. DCVs are delivered from the cell body, and circulate in the neuronal process. Synapsin is required for DCV capture at synapses. In the absence of synapsin and in the S9E mutant, DCVs are not retained at synapses, but accumulate between synapses and in the cell body, likely as they cannot be released. In the S9A mutant, DCVs are anchored to the actin cytoskeleton tightly. Neuropeptide release, based on our electrophysiological recordings, is still possible.

The stationary fluorescent particles may represent synaptic / DCV release sites, near which DCVs are captured and made available for fusion. Distinct capture events, as well as release events could be observed in the kymograms (**Fig. 8b**; see **Supplementary Fig. S3** for all kymograms obtained).

We compared wt and *snn-1(tm2557)* animals, as well as the S9A and S9E mutants. Trafficking of the mobile particles was mostly of uniform velocity, however, S9A and S9E mutants exhibited significantly slower retrograde traffic (**Fig. 8c**). Overall, a similar fraction of the particles observed in each kymograph were stationary, thus, the mutations did not cause a loss or gain of release sites (**Fig. 8d**). However, when we assessed the overall movement for each group of animals, wt showed the highest accumulated distance covered by the mobile particles, while *tm2557* as well as S9E mutants showed significantly less motility. The most prominent reduction was observed for the S9A mutant, lacking the PKA phosphorylation site in the long isoform (SNN-1B; **Fig. 8e**). DCVs in these animals may thus be tightly bound to the actin cytoskeleton. Last, we also analyzed the overall fluorescence. Here, the S9A animals showed a significant (2-fold) increase of DCVs in the axon, compared to wt (**Fig. 8f**), where axon and dendrite showed similar fluorescence levels. Thus S9A appears to cause an accumulation of DCVs in axons. Our findings indicate that SNN-1 phosphorylation by PKA is required to make DCVs mobile and release them from the cytoskeleton. However, also the S9E mutation caused reduced DCV motility, as did the *snn-1(tm2557)* deletion allele, and while these two mutants did not release neuropeptides, the S9A animals did (based on the observed mPSC amplitude increase; **Fig. 6h**). Thus, synapsin (S9 phosphorylation) may affect multiple steps of DCV trafficking and recruitment to release sites, which is differently affected in S9E and S9A mutants.

## DISCUSSION

In this study, we analyzed how synapsin, a known organizer of the SV cluster, functions in *C. elegans* cholinergic motor neurons in fusion of SVs and DCVs and particularly in neuropeptide release. In the *snn-1(tm2557)* deletion mutant, removing the C-terminal half, and likely eliminating SNN-1 expression entirely, we observed some alteration in SV localization. This also caused a mild reduction of SV release, as induced by optogenetic cAMP elevation. However, these effects were small. Instead, we found strong phenotypes and thus a previously unknown role of synapsin in neuropeptide release: *snn-1(tm2557)* mutants, in response to bPAC-induced cAMP increase, showed altered behavior, no neuropeptide release, less overall synaptic DCVs but no acute further reduction of DCVs in terminals, no increase in mPSC amplitudes, and no SV size increase (i.e. ACh loading), as opposed to what we previously found in wt (Steuer Costa et al., 2017). DCV abundance (reduced at synapses, increased in neurites and somata) and localization were affected in the absence of SNN-1, as well as in mutants of the phosphorylation site S9 in SNN-1B: Both abolished (S9A) or ‘constitutive’ phosphorylation (S9E) caused reduced DCV numbers near the center of synapses, while S9A caused a strong accumulation of immobile DCVs in axons, likely because their interaction with the actin cytoskeleton was enhanced. The SNN-1B S9 site may thus affect DCV trafficking and function by regulating their clustering and/or their interaction with the cytoskeleton. At synapses, DCVs may have to be ‘captured’ from their ongoing trafficking along microtubule tracts. However, they also have to be released again to make them available for fusion (see model, **Fig. 8g**). The right sequence of these differential de-/phosphorylation events may be more affected by the synapsin S9E mutation rather than by S9A, because at some point DCVs have to be bound by synapsin to make them available at release site, and this may not happen effectively in S9E mutants.

In the *tm2557* mutants, DCV capture apparently does not occur or is reduced. The lower number of DCVs in *snn-1(tm2557)* synapses showed an altered distribution, and upon stimulation by bPAC, DCVs were depleted near DPs and accumulated distal to the synapse. In SNN-1B(S9A) animals, bPAC stimulation did not alter the DCV distribution at synapses. In S9E animals, like in *tm2557*, bPAC stimulation caused an increase of DCVs distal to the synapse. Possibly, activity causes a redistribution of DCVs but no proper recruitment at release sites occurs, thus DCVs may be ‘dropped’ in these regions of the axon when no synapsin-based tethering of DCVs to the actin cytoskeleton can occur. This supports two conclusions: 1) Contrary to previous publications that attributed ‘capture’ merely to a tightly regulated balance of antero- and retrograde molecular motors of the tubulin cytoskeleton (Bharat et al., 2017; Morrison et al., 2018; Stucchi et al., 2018), our data suggest that also a direct tethering of DCVs occurs by synapsin, as established for SVs, to keep them near synaptic release sites. 2) This process appears to depend on S9, where phosphorylation leads to release of the captured DCVs from the actin cytoskeleton to enable their PM localization and fusion. According to the ‘conveyor belt’ model of DCV delivery to synapses in *Drosophila*, i.e. by circulation in axons and terminals of motor neurons (Wong et al., 2012), also DCVs in *C. elegans* may traffic into the axon, but could then be transported back to the cell body. This is in line with our kymographs of NLP-21∷Venus tagged DCVs. In *Drosophila*, another pre-synaptic PKA target, the RyR, also affects DCV capture at synapses (Wong et al., 2009). In *C. elegans*, UNC-68 RyR was shown to affect SV quantal content in motor neurons (Liu et al., 2005). Whether cAMP effects on DCV release require concomitant depolarization is not clear at present, since the available assays of PSC amplitude depend on SV release, which requires depolarization. The two signaling pathways may be independently used for DCVs (cAMP) and SVs (depolarization).

The *C. elegans* genome encodes one synapsin gene, giving rise to two splice variants, SNN-1A, which lacks the N-terminal A domain and the S9 phosphosite, and SNN-1B, that includes the A domain (**Fig. 1e**). While we have no information of cell-specific or developmentally regulated expression of these two variants, our data indicate that the SNN-1B variant contributes to function in cholinergic neurons. Yet, despite the effects of the *snn-1* deletion on synaptic SV distribution and particularly on SV mobilization from the reserve pool, the bPAC-induced increase of SV release was only partially affected by SNN-1B. Synapsin binds to SV membranes through its central C domain, which is also responsible for dimerization and interaction with the actin cytoskeleton. Phosphorylation of S9 affects the affinity for actin and is responsible for PKA-dependent mobilization of SVs from the RP (Cesca et al., 2010; Hosaka et al., 1999). The *snn-1(tm2557)* deletion eliminates regions for SNN-1 interaction with SVs and actin, and affects dimerization (Esser et al., 1998). Thus, SVs may be less efficiently tethered near synapses. The remaining bPAC-induced increase of mPSC rate in *snn-1(tm2557)* mutants, though lower than in wt, argues that PKA regulation of SV release is not only affected via SNN-1. Hence, fast cAMP effects on SV release likely involve also other PKA targets that affect priming, such as tomosyn (Baba et al., 2005; McEwen et al., 2006) or SNAP-25 (Nagy et al., 2004). Alternatively, SNN-1A, lacking the N-terminal PKA site and A domain, but including all other regions, including the IDR, also mediates some function in cholinergic neurons.

It was shown that ATP acts as a hydrotrope that can dissolve phase-separated IDR-containing proteins (Patel et al., 2017). Might bPAC cause depletion of ATP to a level that could affect synapsin phase-separation (Milovanovic et al., 2018)? We consider this unlikely: First, the concentration of bPAC is likely insufficient to reduce ATP to such levels. Second, even if this was the case, such a drop in ATP levels should affect neuronal function. However, based on the induced behaviors and increased synaptic transmission we observed upon bPAC stimulation, this is not plausible. Third, reducing ATP levels should actually increase phase separation and thus counteract cAMP-induced SV mobilization and increase of synaptic transmission.

SNN-1 is required for DCV fusion. Discharged neuropeptides activate auto-receptors, increasing the ACh load of SVs (Steuer Costa et al., 2017). Likewise, without UNC-31/CAPS, no neuropeptide release occurs, eliminating effects on mPSC amplitude, but not their rate. CAPS-dependent signaling in secretory granule filling was found in chromaffin cells of mice, as CAPS1 knockouts were deficient in catecholamine-loading (Speidel et al., 2005) and showed reduced DCV priming and release (Liu et al., 2008). CAPS1 and 2 were also shown to be required for SV priming in mice (Jockusch et al., 2007). This may be reflected in *C. elegans*, as *unc-31* mutants lacked not only the cAMP-induced amplitude increase but also had a reduced mPSC rate (Steuer Costa et al., 2017). Unlike the findings of Charlie et al. (Charlie et al., 2006a), our data place UNC-31 downstream of cAMP/PKA effects on synapsin, and upstream of neuropeptide release in evoking increased SV quantal size. Thus, the unknown PKA target postulated by Zhou et al. (Zhou et al., 2007) likely is SNN-1. cAMP signaling would enable synapses to undergo homeostatic changes, in response to altered signaling at the NMJ, possibly to adapt to different locomotion regimes. cAMP-dependent synaptic facilitation changes behavior within seconds to tens of seconds. The intrinsic upstream trigger for cAMP signaling at cholinergic synapses remains to be identified. Recently, neuromodulators were shown to affect synapsin phosphorylation after 30 min through cAMP pathways, and to alter SV numbers in murine and human neurons (Patzke et al., 2019). PKA inhibition increased, and cAMP upregulation decreased SV numbers. Though the authors did not assess rapid effects after a few seconds of cAMP increase, the observed decrease in SV numbers indicates that cAMP/SNN-1 pathways of SV regulation are conserved during evolution. These authors also analyzed a number of mutations of synapsin found in human genetic diseases. Our work implies that these pathologies may, at least in part, be mediated at the level of neuropeptide signaling.

---

**Fig. S1:**
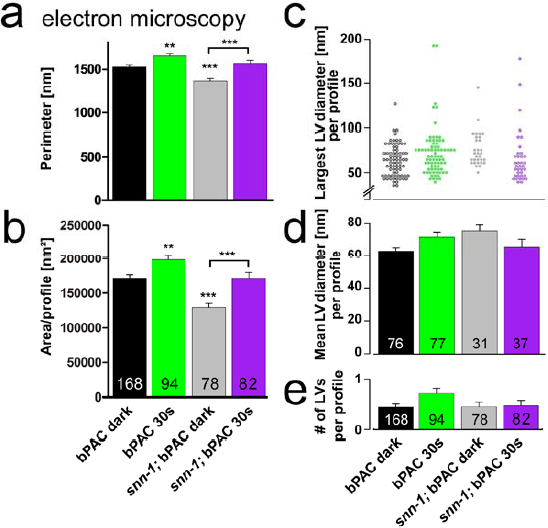
Morphological characteristics of synapses in wt and *snn-1(tm2557)* mutants, as well as number and size of bPAC-stimulation induced large vesicles. **a, b)** Mean size characteristics of synapses analyzed by EM in this work. Mean perimeter and area per profile, per condition and genotype as indicated. **c-e)** Quantification of endocytic large vesicles (LVs), induced by bPAC stimulation. **c)** Largest diameter of individual LVs observed, **d)** mean LV diameter per synaptic profile, **e)** mean LV number per synaptic profile. *p≤0.05, **p≤0.01, ***≤0.001, one-way ANOVA with Tukey correction.

**Fig. S2:**
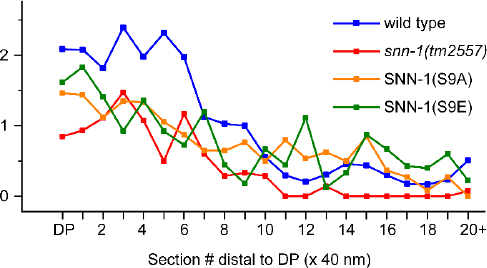
SNN-1B deletion and serine 9 mutations affect DCV distribution in sections containing dense projections and flanking sections. As in Fig. 7, but comparing mean DCV distribution in indicated genotypes in the non-stimulated condition. SEM not shown for clarity (see Fig. 7).

**Fig. S3:**
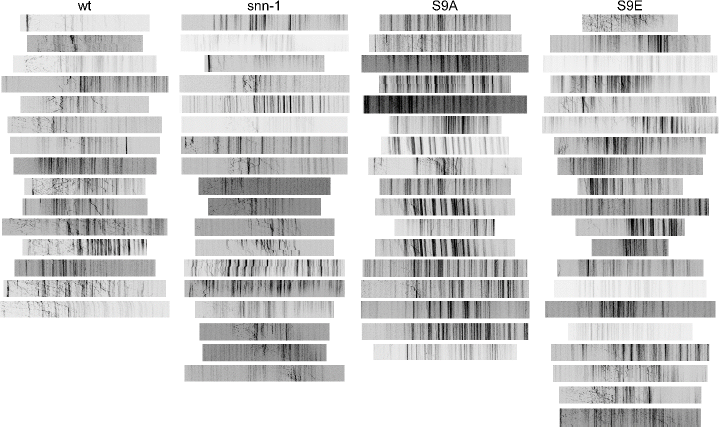
DCV kymograms of all analyzed animals, sorted by genotype, related to Fig. 8.

## AUTHOR CONTRIBUTIONS

Experiments were conceived, designed, and analyzed by S.-c. Y., W.S.C., J.F.L., J.S. and A.G. The manuscript was written by A.G. with help by the other authors.

## ACKNOWLEDGMENTS

We are grateful to K. Miller and J. Kaplan for providing strains, and to Mona Hoeret and Franziska Baumbach for expert technical assistance. Some nematode strains were obtained from the *Caenorhabditis* Genetics Center (CGC) which is funded by the NIH National Center for Research Resources (NCRR), and from the Japanese National Bioresource Project for the Experimental Animal “Nematode *C. elegans*”. This work was funded by the Deutsche Forschungsgemeinschaft (DFG), grants CRC1080-B2, GO1011/4−1/−2, and EXC115, and was further supported by Goethe University.

## EXPERIMENTAL PROCEDURES

### Strains and Genetics

*C. elegans* strains were cultivated using standard methods on nematode growth medium (NGM) and fed *E. coli* strain OP50-1 (Brenner, 1974). For all behavioral and electrophysiology experiments, animals were in *lite-1(ce314)* background (lacking the intrinsic photophobic response) (Edwards et al., 2008), for all bPAC EM work, the wild type background was used. Strains used or generated: *snn-1(tm2557)*, **KG1034:***ceIs33[rab-3∷Tpde-4d(+) cDNA]*, **KG1180:***lite-1(ce314)*, **RB830:***epac-1(ok655)*, **TR2171:***unc-68(r1162)*, **ZX1460:***ceIs33[rab-3∷Tpde-4d(+) cDNA]; zxIs53[punc-17∷bPAC∷YFP, pmyo-2∷mCherry]*, **ZX1461:***zxIs53[punc-17∷bPAC∷YFP, pmyo-2∷mCherry]*, **ZX1555:** *nuIs183[Punc-129∷NLP-21∷Venus; myo-2∷NLS∷GFP]*, **ZX1569:***lite-1(ce314); zxIs53[punc-17∷bPAC∷YFP; pmyo-2∷mCherry]*, **ZX1570:***lite-1(ce314); unc-31(n1304); zxIs53[punc-17∷bPAC∷YFP; pmyo-2∷mCherry]*, **ZX1867:***epac-1(ok655); zxIs53[punc-17∷bPAC∷YFP; pmyo-2∷mCherry]*, **ZX1868:***snn-1(tm2557); lite-1(ce314)*, **ZX1870:***unc-31(n1304); zxIs53[punc-17∷bPAC∷YFP; pmyo-2∷mCherry]*, **ZX1871:***snn-1(tm2557); lite-1(ce314); zxIs53[punc-17∷bPAC∷YFP; pmyo-2∷mCherry]*, **ZX1990:***lite-1(ce314); zxIs53[punc-17∷bPAC∷YFP; pmyo-2∷mCherry] nuIs183[Punc-129∷NLP-21∷Venus; myo-2∷NLS∷GFP]*, **ZX1991:***snn-1(tm2557); lite-1(ce314); zxIs53[punc-17∷bPAC∷YFP; pmyo-2∷mCherry] nuIs183[punc-129∷NLP-21∷Venus; pmyo-2∷NLS∷GFP]*, **ZX1992:***unc-68(r1162), zxIs53[punc-17∷bPAC∷YFP; pmyo-2∷mCherry]*, **ZX2553:***snn-1(tm2557); nuIs183[punc-129∷NLP-21∷Venus; pmyo-2∷NLS∷GFP]*, **ZX2555:***snn-1(S9E); nuIs183[punc-129∷NLP-21∷Venus; pmyo-2∷NLS∷GFP]*, **ZX2559:** *snn-1(S9A); nuIs183[punc-129∷NLP-21∷Venus; pmyo-2∷NLS∷GFP]*.

### Molecular biology and CRISPR/Cas9 genome editing

The construct for the photoactivated adenylyl cyclase of *Beggiatoa sp.* was previously described (Steuer Costa et al., 2017). The *snn-1(S9A)* and *snn-1(S9E)* point mutations were engineered by CRISPR/Cas9 genome editing.

### Generation of transgenic animals

Generation of transgenic *C. elegans* expressing punc-17∷bPAC∷YFP; pmyo-2∷mCherry (strain ZX1460) was previously described (Steuer Costa et al., 2017). This transgene, *zxIs53*, was used for crossing into the mutant alleles tested in this work.

### Behavioral assays

Locomotion parameters on NGM were acquired with a previously described worm tracker (Stirman et al., 2011) with the following modifications: First, a mechanical shutter (Sutter Instrument Company, Novato, CA 94949, USA) was placed between projector and microscope and synchronized to the light stimulation. Second, a band pass filter (650 ± 25 nm) was inserted in the background light path. These modifications ensured an ambient light power, measured between 200 nm and 1000 nm, below 20 nW/mm^2^ during tracking in ‘dark’. Light power was measured with a powermeter (PM100, Thorlabs, Newton, NY, USA) at the focal plane while the sensor was placed at the expected worm’s position. All animals were kept in darkness during growth. Young adults were transferred individually on plain NGM plates under red light (>600 nm) in a dark room and kept for 15 minutes in the dark before transfer to the tracker. Light stimulation protocol was 15 s in darkness, 25 s in 70 μW/mm^2^ 470/10 nm light and 15 s darkness. Tracks were automatically filtered to exclude data points from erroneously evaluated movie frames with a custom workflow in KNIME (KNIME Desktop version 2.10, KNIME.com AG, 8005 Zurich, Switzerland) (Preisach et al., 2008; Warr, 2012). Our constraints were that animals do not move faster than 1.25 mm/s and their length does not show a discrepancy above 25% to the mean first five seconds of the video. Videos were excluded from analysis when more than 15% of the data points had to be discarded by our constraints. Behavior data passed the Shapiro-Wilk normality test. Bending angle analysis during crawling was defined by the mean deviation from 180° from 11 equally interspaced 3-point angles defined by 13 points along the spine of the animal.Behavior data clustering was performed on the absolute change compared to the mean value before light using dynamic time warping distances and the hclust function with method “average” in R (Version 3.2.1, libraries dtw, ape, gplots; http://www.R-project.org/).

### Electrophysiology

Recordings from dissected body wall muscle cells were conducted as described previously (Liewald et al., 2008). Light activation was performed using an LED lamp (KSL-70, Rapp OptoElectronic, Hamburg, Germany; 470 nm, 8 mW/mm^2^) and controlled by an EPC10 amplifier and Patchmaster software (HEKA, Germany). mPSC analysis was done by Mini Analysis software (Synaptosoft, Decatur, GA, USA, version 6.0.7). Amplitude and mean number of mPSC events per second were analyzed during the following time bins: 30 s before illumination, the first 25 s of illumination and the last 30 s after illumination.

### Electron Microscopy

Young adult animals were used for HPF fixation, based on methods previously described (Weimer, 2006). Briefly, about 10-20 worms were loaded into a 100 μm deep aluminum planchette (Microscopy Services) filled with *E. coli* OP50, covered with a 0.16 mm sapphire disc and a 0.4 mm spacer ring (Engineering office M. Wohlwend GmbH) for subsequent photostimulation. To prevent preactivation of bPAC, all manipulations were done under low-light conditions or under red light. For light stimulation experiments, worms were continuously illuminated with a ~470nm blue LED (3 mW/mm^2^) for 30s, followed by high-pressure freezing at −180°C under 2,100 bar pressure in a Bal-Tec HPM010 or a Leica HPM100 machine. After freezing, specimens were transferred under liquid nitrogen into a Reichert AFS machine (Leica) for freeze substitution. Tannic acid (0.1% in dry acetone) fixative was used to incubate samples at −90°C for 100 hours. Then, a process of washing was performed to substitute with acetone, followed by an incubation of 2% OsO_4_ for 39.5 hours (in dry acetone) while slowly increasing temperature up to room temperature. Afterwards, Epoxy resin (Agar Scientific, AGAR 100 Premix kit hard) embedding process was executed with increasing concentration from 50% to 90% at room temperature and 100% at 60°C over 48 hours. Cross sections were cut at a thickness of 40 nm, transferred on formvar-covered copper slot grids and counterstained in 2.5% aqueous uranyl acetate for 4 min, followed by distilled water wash. Then, grids were carried onto Reynolds lead citrate solution for 2 min in a CO_2_-free chamber and then washed in distilled water again. Images of regions in the ventral nerve cord were taken with a Erlangshen ES500W CCD camera (Gatan) in a Philips CM12 operated at 80kV. Images were tagged in ImageJ (1.47v, NIH) for synapse perimeter, SVs, docked SVs, DCVs, LVs, and dense projection, scored blind for each condition. ImageJ ROIs were stored for position, area, circumference and largest diameter, and then quantified and automatically analyzed by custom software written in R, called by a KNIME workflow. The diameters of synapses from each stimulation condition varied in size, thus, each value for the number of docked SVs was normalized to the mean perimeter and represents the number of docked SVs along a membrane whose perimeter is 1,548 nm in a profile. The other organelles are represented as the numbers of SVs, DCVs or LVs in a typical synaptic profile of 164,100 nm^2^. 3D reconstructions of serial sections were performed as described earlier (Kittelmann et al., 2013; Steuer Costa et al., 2017).

### Fluorescence imaging

For coelomocyte imaging, animals were either kept in dark or illuminated for 15 min with a 470 nm LED, 350 μW/mm^2^ at 20°C. Images were acquired up to 30 min after illumination. Image acquisition was performed with a Zeiss Cell Observer SD with an alpha Plan-Apochromoat 100×/1.46 Oil DIC (UV) objective, 488 nm excitation laser at 40% power and a Rolera EM-C2 with EM gain of 100, full resolution and 100 ms exposure time. Images were exported as 16 bit .tif files. In ImageJ (1.47v, NIH), ROIs were traced for the anterior coelomocyte and a background area outside of the animal. Similarly, cell bodies and processes in the ventral nerve cord were analyzed for fluorescence. Mean intensity of these ROIs were exported and fluorescence intensity was corrected for background intensity prior to normalization to the NLP-21∷Venus marker strain in the dark (Sieburth et al., 2007) (Venus is an enhanced YFP variant).

### Analysis of trafficking of NLP-21∷Venus containing DCVs

z-stacks of fluorescence images, enclosing the dorsal nerve cord, were acquired on a Zeiss Cell Observer Spinning Disk confocal microscope with a Plan-Apochromat 63×/1.40 Oil DIC M27 objective (Zeiss), 488 nm excitation laser at 20% power and a Rolera EM-C2 EMCCD camera with EM gain of 100, full resolution and 70 ms exposure time. Live animals were immobilized on a 10% agarose pad (in M9 buffer) supplemented with a drop of 20 mM tetramisole, under a coverslip. To generate kymographs, we used Image J and the ‘Velocity Measurement Tool’ (http://dev.mri.cnrs.fr/projects/imagej-macros/wiki/Velocity_Measurement_Tool). A segmented line was drawn along the nerve cord process (i.e. the path of moving particles) and used to obtain line scan kymographs. Trajectories of single DCVs for each kymograph were traced and analyzed for antero- or retrograde trafficking, and velocity, by using the respective features of the macro. Further analysis was done using Microsoft Excel.

### Statistics

Data is given as means ± SEM. A summary of acquired EM data and statistics analysis is given in the **Supplementary Table 1**. Significance between data sets after two-tailed Student’s t-test or after ANOVA is given as p-value (* p ≤ 0.05; ** p ≤ 0.01; *** p ≤ 0.001), the latter after Bonferroni’s multiple comparison test, or Tukey’s post-hoc test. Data was analyzed and plotted in GraphPad Prism (GraphPad Software, Inc., La Jolla, CA, USA’, version 5.01), or R (Version 3.2.1, libraries dtw, ape, gplots; http://www.R-project.org/), or in OriginPro 2015G (OriginLab Corporation, Northampton, USA). For empirical cumulative distribution functions (eCDFs), the sum of SV linear distances to the DP was divided by the profile area, to represent the distribution of SVs relative to the DP in a profile area-independent manner. These values were plotted as an eCDF to show the distribution of SVs across the profiles in one group. Equivalency of distributions was analyzed by a Kolmogorov-Smirnov test (KS-Test) between groups. We report the empirical distribution functions as originating from different distribution functions when the KS-Test p-value is smaller than 0.05. We conclude that the distribution of SVs relative to the DP is changed across groups that fulfill this statement.

## SUPPLEMENTARY MATERIAL

**Supplementary Table S1** Statistical analysis of electron microscopy data presented in this work

**Supplementary Video S1** Trafficking of NLP-21∷Venus containing DCVs in the dendrite of a DA type motor neuron.

## REFERENCES

Baba, T., Sakisaka, T., Mochida, S., and Takai, Y. (2005). PKA-catalyzed phosphorylation of tomosyn and its implication in Ca2+-dependent exocytosis of neurotransmitter. J Cell Biol 170, 1113–1125.

Bark, S.J., Wegrzyn, J., Taupenot, L., Ziegler, M., O’Connor, D.T., Ma, Q., Smoot, M., Ideker, T., and Hook, V. (2012). The protein architecture of human secretory vesicles reveals differential regulation of signaling molecule secretion by protein kinases. PLoS One 7, e41134.

Benfenati, F., Bahler, M., Jahn, R., and Greengard, P. (1989a). Interactions of synapsin I with small synaptic vesicles: distinct sites in synapsin I bind to vesicle phospholipids and vesicle proteins. J Cell Biol 108, 1863–1872.

Benfenati, F., Valtorta, F., Bahler, M., and Greengard, P. (1989b). Synapsin I, a neuron-specific phosphoprotein interacting with small synaptic vesicles and F-actin. Cell biology international reports 13, 1007–1021.

Bharat, V., Siebrecht, M., Burk, K., Ahmed, S., Reissner, C., Kohansal-Nodehi, M., Steubler, V., Zweckstetter, M., Ting, J.T., and Dean, C. (2017). Capture of Dense Core Vesicles at Synapses by JNK-Dependent Phosphorylation of Synaptotagmin-4. Cell reports 21, 2118–2133.

Brenner, S. (1974). The genetics of Caenorhabditis elegans. Genetics 77, 71–94.

Burkhardt, P. (2015). The origin and evolution of synaptic proteins-choanoflagellates lead the way. J Exp Biol 218, 506–514.

Cavolo, S.L., Bulgari, D., Deitcher, D.L., and Levitan, E.S. (2016). Activity Induces Fmr1-Sensitive Synaptic Capture of Anterograde Circulating Neuropeptide Vesicles. J Neurosci 36, 11781–11787.

Cesca, F., Baldelli, P., Valtorta, F., and Benfenati, F. (2010). The synapsins: Key actors of synapse function and plasticity. Progress in Neurobiology 91, 313–348.

Charlie, N.K., Schade, M.A., Thomure, A.M., and Miller, K.G. (2006a). Presynaptic UNC-31 (CAPS) is required to activate the G alpha(s) pathway of the Caenorhabditis elegans synaptic signaling network. Genetics 172, 943–961.

Charlie, N.K., Thomure, A.M., Schade, M.A., and Miller, K.G. (2006b). The Dunce cAMP phosphodiesterase PDE-4 negatively regulates G alpha(s)-dependent and G alpha(s)-independent cAMP pools in the Caenorhabditis elegans synaptic signaling network. Genetics 173, 111–130.

Cheung, U., Atwood, H.L., and Zucker, R.S. (2006). Presynaptic effectors contributing to cAMP-induced synaptic potentiation in Drosophila. J Neurobiol 66, 273–280.

Cho, R.W., Buhl, L.K., Volfson, D., Tran, A., Li, F., Akbergenova, Y., and Littleton, J.T. (2015). Phosphorylation of Complexin by PKA Regulates Activity-Dependent Spontaneous Neurotransmitter Release and Structural Synaptic Plasticity. Neuron 88, 749–761.

Edwards, S.L., Charlie, N.K., Milfort, M.C., Brown, B.S., Gravlin, C.N., Knecht, J.E., and Miller, K.G. (2008). A novel molecular solution for ultraviolet light detection in Caenorhabditis elegans. PLoS Biol 6, 0060198.

Esser, L., Wang, C.R., Hosaka, M., Smagula, C.S., Sudhof, T.C., and Deisenhofer, J. (1998). Synapsin I is structurally similar to ATP-utilizing enzymes. EMBO J 17, 977–984.

Evans, G.J., Wilkinson, M.C., Graham, M.E., Turner, K.M., Chamberlain, L.H., Burgoyne, R.D., and Morgan, A. (2001). Phosphorylation of cysteine string protein by protein kinase A. Implications for the modulation of exocytosis. J Biol Chem 276, 47877–47885.

Gekel, I., and Neher, E. (2008). Application of an Epac activator enhances neurotransmitter release at excitatory central synapses. J Neurosci 28, 7991–8002.

Gitler, D., Cheng, Q., Greengard, P., and Augustine, G.J. (2008). Synapsin IIa controls the reserve pool of glutamatergic synaptic vesicles. J Neurosci 28, 10835–10843.

Gracheva, E.O., Burdina, A.O., Holgado, A.M., Berthelot-Grosjean, M., Ackley, B.D., Hadwiger, G., Nonet, M.L., Weimer, R.M., and Richmond, J.E. (2006). Tomosyn inhibits synaptic vesicle priming in Caenorhabditis elegans. PLoS Biol 4, e261.

Graebner, A.K., Iyer, M., and Carter, M.E. (2015). Understanding how discrete populations of hypothalamic neurons orchestrate complicated behavioral states. Frontiers in systems neuroscience 9, 111.

Hammarlund, M., Watanabe, S., Schuske, K., and Jorgensen, E.M. (2008). CAPS and syntaxin dock dense core vesicles to the plasma membrane in neurons. J Cell Biol 180, 483–491.

Haucke, V., Neher, E., and Sigrist, S.J. (2011). Protein scaffolds in the coupling of synaptic exocytosis and endocytosis. Nat Rev Neurosci 12, 127–138.

Hoover, C.M., Edwards, S.L., Yu, S.C., Kittelmann, M., Richmond, J.E., Eimer, S., Yorks, R.M., and Miller, K.G. (2014). A novel CaM kinase II pathway controls the location of neuropeptide release from Caenorhabditis elegans motor neurons. Genetics 196, 745–765.

Hosaka, M., Hammer, R.E., and Sudhof, T.C. (1999). A phospho-switch controls the dynamic association of synapsins with synaptic vesicles. Neuron 24, 377–387.

Hua, Y., Woehler, A., Kahms, M., Haucke, V., Neher, E., and Klingauf, J. (2013). Blocking endocytosis enhances short-term synaptic depression under conditions of normal availability of vesicles. Neuron 80, 343–349.

Jahn, R., and Fasshauer, D. (2012). Molecular machines governing exocytosis of synaptic vesicles. Nature 490, 201–207.

Jahne, S., Rizzoli, S.O., and Helm, M.S. (2015). The structure and function of presynaptic endosomes. Exp Cell Res 335, 172–179.

Jockusch, W.J., Speidel, D., Sigler, A., Sorensen, J.B., Varoqueaux, F., Rhee, J.S., and Brose, N. (2007). CAPS-1 and CAPS-2 are essential synaptic vesicle priming proteins. Cell 131, 796–808.

Johnson, E.M., Ueda, T., Maeno, H., and Greengard, P. (1972). Adenosine 3’,5-monophosphate-dependent phosphorylation of a specific protein in synaptic membrane fractions from rat cerebrum. J Biol Chem 247, 5650–5652.

Kaneko, M., and Takahashi, T. (2004). Presynaptic mechanism underlying cAMP-dependent synaptic potentiation. J Neurosci 24, 5202–5208.

Kittelmann, M., Liewald, J.F., Hegermann, J., Schultheis, C., Brauner, M., Steuer Costa, W., Wabnig, S., Eimer, S., and Gottschalk, A. (2013). In vivo synaptic recovery following optogenetic hyperstimulation. Proc Natl Acad Sci U S A 110, E3007–3016.

Kononenko, N.L., and Haucke, V. (2015). Molecular mechanisms of presynaptic membrane retrieval and synaptic vesicle reformation. Neuron 85, 484–496.

Kuromi, H., and Kidokoro, Y. (2000). Tetanic stimulation recruits vesicles from reserve pool via a cAMP-mediated process in Drosophila synapses. Neuron 27, 133–143.

Liewald, J.F., Brauner, M., Stephens, G.J., Bouhours, M., Schultheis, C., Zhen, M., and Gottschalk, A. (2008). Optogenetic analysis of synaptic function. Nat Methods 5, 895–902.

Liu, Q., Chen, B., Yankova, M., Morest, D.K., Maryon, E., Hand, A.R., Nonet, M.L., and Wang, Z.W. (2005). Presynaptic ryanodine receptors are required for normal quantal size at the Caenorhabditis elegans neuromuscular junction. J Neurosci 25, 6745–6754.

Liu, Y., Schirra, C., Stevens, D.R., Matti, U., Speidel, D., Hof, D., Bruns, D., Brose, N., and Rettig, J. (2008). CAPS facilitates filling of the rapidly releasable pool of large dense-core vesicles. J Neurosci 28, 5594–5601.

Lonart, G., Schoch, S., Kaeser, P.S., Larkin, C.J., Sudhof, T.C., and Linden, D.J. (2003). Phosphorylation of RIM1alpha by PKA triggers presynaptic long-term potentiation at cerebellar parallel fiber synapses. Cell 115, 49–60.

McEwen, J.M., Madison, J.M., Dybbs, M., and Kaplan, J.M. (2006). Antagonistic Regulation of Synaptic Vesicle Priming by Tomosyn and UNC-13. Neuron 2006, 303–315.

Menegon, A., Bonanomi, D., Albertinazzi, C., Lotti, F., Ferrari, G., Kao, H.T., Benfenati, F., Baldelli, P., and Valtorta, F. (2006). Protein kinase A-mediated synapsin I phosphorylation is a central modulator of Ca2+-dependent synaptic activity. J Neurosci 26, 11670–11681.

Milovanovic, D., Wu, Y., Bian, X., and De Camilli, P. (2018). A liquid phase of synapsin and lipid vesicles. Science 361, 604–607.

Morrison, L.M., Edwards, S.L., Manning, L., Stec, N., Richmond, J.E., and Miller, K.G. (2018). Sentryn and SAD Kinase Link the Guided Transport and Capture of Dense Core Vesicles in Caenorhabditis elegans. Genetics 210, 925–946.

Nagy, G., Reim, K., Matti, U., Brose, N., Binz, T., Rettig, J., Neher, E., and Sorensen, J.B. (2004). Regulation of releasable vesicle pool sizes by protein kinase A-dependent phosphorylation of SNAP-25. Neuron 41, 417–429.

Oranth, A., Schultheis, C., Tolstenkov, O., Erbguth, K., Nagpal, J., Hain, D., Brauner, M., Wabnig, S., Steuer Costa, W., McWhirter, R.D., et al. (2018). Food Sensation Modulates Locomotion by Dopamine and Neuropeptide Signaling in a Distributed Neuronal Network. Neuron 100, 1414–1428 e1410.

Park, Y.S., Hur, E.M., Choi, B.H., Kwak, E., Jun, D.J., Park, S.J., and Kim, K.T. (2006). Involvement of protein kinase C-epsilon in activity-dependent potentiation of large dense-core vesicle exocytosis in chromaffin cells. J Neurosci 26, 8999–9005.

Patel, A., Malinovska, L., Saha, S., Wang, J., Alberti, S., Krishnan, Y., and Hyman, A.A. (2017). ATP as a biological hydrotrope. Science 356, 753–756.

Patzke, C., Brockmann, M.M., Dai, J., Gan, K.J., Grauel, M.K., Fenske, P., Liu, Y., Acuna, C., Rosenmund, C., and Sudhof, T.C. (2019). Neuromodulator Signaling Bidirectionally Controls Vesicle Numbers in Human Synapses. Cell 179, 498–513 e422.

Preisach, C., Burkhardt, H., Schmidt-Thieme, L., and Decker, R., eds. (2008). Data Analysis, Machine Learning and Applications (Berlin; Heidelberg: Springer Berlin Heidelberg).

Rizzoli, S.O. (2014). Synaptic vesicle recycling: steps and principles. EMBO J 33, 788–822.

Rizzoli, S.O., and Betz, W.J. 2005). Synaptic vesicle pools. Nat Rev Neurosci 6, 57–69.

Rodriguez, P., Bhogal, M.S., and Colyer, J. (2003). Stoichiometric phosphorylation of cardiac ryanodine receptor on serine 2809 by calmodulin-dependent kinase II and protein kinase A. J Biol Chem 278, 38593–38600.

Rupnik, M., Kreft, M., Sikdar, S.K., Grilc, S., Romih, R., Zupancic, G., Martin, T.F., and Zorec, R. (2000). Rapid regulated dense-core vesicle exocytosis requires the CAPS protein. Proc Natl Acad Sci U S A 97, 5627–5632.

Schneggenburger, R., and Rosenmund, C. (2015). Molecular mechanisms governing Ca(2+) regulation of evoked and spontaneous release. Nat Neurosci 18, 935–941.

Sieburth, D., Madison, J.M., and Kaplan, J.M. (2007). PKC-1 regulates secretion of neuropeptides. Nat Neurosci 10, 49–57.

Sigrist, S.J., and Schmitz, D. (2011). Structural and functional plasticity of the cytoplasmic active zone. Curr Opin Neurobiol 21, 144–150.

Siksou, L., Rostaing, P., Lechaire, J.P., Boudier, T., Ohtsuka, T., Fejtova, A., Kao, H.T., Greengard, P., Gundelfinger, E.D., Triller, A., et al. (2007). Three-dimensional architecture of presynaptic terminal cytomatrix. J Neurosci 27, 6868–6877.

Speese, S., Petrie, M., Schuske, K., Ailion, M., Ann, K., Iwasaki, K., Jorgensen, E.M., and Martin, T.F. (2007). UNC-31 (CAPS) is required for dense-core vesicle but not synaptic vesicle exocytosis in Caenorhabditis elegans. J Neurosci 27, 6150–6162.

Speidel, D., Bruederle, C.E., Enk, C., Voets, T., Varoqueaux, F., Reim, K., Becherer, U., Fornai, F., Ruggieri, S., Holighaus, Y., et al. (2005). CAPS1 regulates catecholamine loading of large dense-core vesicles. Neuron 46, 75–88.

Steuer Costa, W., Van der Auwera, P., Glock, C., Liewald, J.F., Bach, M., Schuler, C., Wabnig, S., Oranth, A., Masurat, F., Bringmann, H., et al. (2019). A GABAergic and peptidergic sleep neuron as a locomotion stop neuron with compartmentalized Ca2+ dynamics. Nature communications 10, 4095.

Steuer Costa, W., Yu, S.C., Liewald, J.F., and Gottschalk, A. (2017). Fast cAMP Modulation of Neurotransmission via Neuropeptide Signals and Vesicle Loading. Curr Biol 27, 495–507.

Stigloher, C., Zhan, H., Zhen, M., Richmond, J., and Bessereau, J.L. (2011). The presynaptic dense projection of the Caenorhabditis elegans cholinergic neuromuscular junction localizes synaptic vesicles at the active zone through SYD-2/liprin and UNC-10/RIM-dependent interactions. J Neurosci 31, 4388–4396.

Stirman, J.N., Crane, M.M., Husson, S.J., Wabnig, S., Schultheis, C., Gottschalk, A., and Lu, H. (2011). Real-time multimodal optical control of neurons and muscles in freely behaving Caenorhabditis elegans. Nat Methods 8, 153–158.

Stucchi, R., Plucinska, G., Hummel, J.J.A., Zahavi, E.E., Guerra San Juan, I., Klykov, O., Scheltema, R.A., Altelaar, A.F.M., and Hoogenraad, C.C. (2018). Regulation of KIF1A-Driven Dense Core Vesicle Transport: Ca(2+)/CaM Controls DCV Binding and Liprin-alpha/TANC2 Recruits DCVs to Postsynaptic Sites. Cell reports 24, 685–700.

Sudhof, T.C. (2012). The presynaptic active zone. Neuron 75, 11–25.

Sudhof, T.C. (2013). Neurotransmitter release: the last millisecond in the life of a synaptic vesicle. Neuron 80, 675–690.

Sudhof, T.C., Czernik, A.J., Kao, H.T., Takei, K., Johnston, P.A., Horiuchi, A., Kanazir, S.D., Wagner, M.A., Perin, M.S., De Camilli, P., et al. (1989). Synapsins: mosaics of shared and individual domains in a family of synaptic vesicle phosphoproteins. Science 245, 1474–1480.

Taghert, P.H., and Nitabach, M.N. (2012). Peptide neuromodulation in invertebrate model systems. Neuron 76, 82–97.

Thakur, P., Stevens, D.R., Sheng, Z.H., and Rettig, J. (2004). Effects of PKA-mediated phosphorylation of Snapin on synaptic transmission in cultured hippocampal neurons. J Neurosci 24, 6476–6481.

Wang, H., and Sieburth, D. (2013). PKA controls calcium influx into motor neurons during a rhythmic behavior. PLoS Genet 9, e1003831.

Warr, W.A. (2012). Scientific workflow systems: Pipeline Pilot and KNIME. J Comput Aided Mol Des 26, 801–804.

Watanabe, S., Liu, Q., Davis, M.W., Hollopeter, G., Thomas, N., Jorgensen, N.B., and Jorgensen, E.M. (2013a). Ultrafast endocytosis at Caenorhabditis elegans neuromuscular junctions. eLife 2, e00723.

Watanabe, S., Rost, B.R., Camacho-Perez, M., Davis, M.W., Sohl-Kielczynski, B., Rosenmund, C., and Jorgensen, E.M. (2013b). Ultrafast endocytosis at mouse hippocampal synapses. Nature 504, 242–247.

Watanabe, S., Trimbuch, T., Camacho-Perez, M., Rost, B.R., Brokowski, B., Sohl-Kielczynski, B., Felies, A., Davis, M.W., Rosenmund, C., and Jorgensen, E.M. (2014). Clathrin regenerates synaptic vesicles from endosomes. Nature 515, 228–233.

Weimer, R.M. (2006). Preservation of C. elegans tissue via high-pressure freezing and freeze-substitution for ultrastructural analysis and immunocytochemistry. Methods Mol Biol 351, 203–221.

Wierda, K.D., Toonen, R.F., de Wit, H., Brussaard, A.B., and Verhage, M. (2007). Interdependence of PKC-dependent and PKC-independent pathways for presynaptic plasticity. Neuron 54, 275–290.

Wong, M.Y., Shakiryanova, D., and Levitan, E.S. (2009). Presynaptic ryanodine receptor-CamKII signaling is required for activity-dependent capture of transiting vesicles. J Mol Neurosci 37, 146–150.

Wong, M.Y., Zhou, C., Shakiryanova, D., Lloyd, T.E., Deitcher, D.L., and Levitan, E.S. (2012). Neuropeptide delivery to synapses by long-range vesicle circulation and sporadic capture. Cell 148, 1029–1038.

Xue, L., and Wu, L.G. (2010). Post-tetanic potentiation is caused by two signalling mechanisms affecting quantal size and quantal content. J Physiol 588, 4987–4994.

Yu, S.C., Janosi, B., Liewald, J.F., Wabnig, S., and Gottschalk, A. (2018). Endophilin A and B Join Forces With Clathrin to Mediate Synaptic Vesicle Recycling in Caenorhabditis elegans. Frontiers in molecular neuroscience 11, 196.

Zhong, N., and Zucker, R.S. (2005). cAMP acts on exchange protein activated by cAMP/cAMP-regulated guanine nucleotide exchange protein to regulate transmitter release at the crayfish neuromuscular junction. J Neurosci 25, 208–214.

Zhou, K.M., Dong, Y.M., Ge, Q., Zhu, D., Zhou, W., Lin, X.G., Liang, T., Wu, Z.X., and Xu, T. (2007). PKA activation bypasses the requirement for UNC-31 in the docking of dense core vesicles from C. elegans neurons. Neuron 56, 657–669.

